# JZL184 increases anxiety-like behavior and does not reduce alcohol consumption in female rats after repeated mild traumatic brain injury

**DOI:** 10.1101/2023.05.30.542943

**Authors:** Alejandra Jacotte-Simancas, Patricia Molina, Nicholas Gilpin

**Affiliations:** Department of Physiology, Louisiana State University Health Sciences Center, New Orleans, LA; Alcohol and Drug of Abuse Center of Excellence, LSUHSC, New Orleans, LA; Department of Cell Biology and Anatomy, LSUHSC, New Orleans, LA; Southeast Louisiana VA Healthcare System, New Orleans, LA

## Abstract

Alcohol use disorder (AUD) is highly comorbid with traumatic brain injury (TBI). Previously, using a lateral fluid percussion model (LFP) (an open model of head injury) to generate a single mild to moderate traumatic brain injury (TBI), we showed that TBI produces escalation in alcohol drinking, that alcohol exposure negatively impacts TBI outcomes, and that the endocannabinoid degradation inhibitor (JZL184) confers significant protection from behavioral and neuropathological outcomes in male rodents. In the present study, we used a weight drop model (a closed model of head injury) to produce a repeated mild TBI (rmTBI, 3 TBIs, spaced by 24 hours) to examine the sex-specific effects on alcohol consumption and anxiety-like behavior in rats, and whether systemic treatment with JZL184 would reverse TBI effects on those behaviors in both sexes. In two separate studies, adult male and female Wistar rats were subjected to rmTBI or sham using the weight drop model. Physiological measures of injury severity were collected from all animals. Animals in both studies were allowed to consume alcohol using an intermittent 2-bottle choice procedure (12 pre-TBI sessions and 12 post-TBI sessions). Neurological severity and neurobehavioral scores (NSS and NBS, respectively) were tested 24 hours after the final injury. Anxiety-like behavior was tested at 37-38 days post-injury in Study 1, and 6-8 days post-injury in Study 2. Our results show that females exhibited reduced respiratory rates relative to males with no significant differences between Sham and rmTBI, no effect of rmTBI or sex on righting reflex, and increased neurological deficits in rmTBI groups in both studies. In Study 1, rmTBI increased alcohol consumption in female but not male rats. Male rats consistently exhibited higher levels of anxiety-like behavior than females. rmTBI did not affect anxiety-like behavior 37-38 days post-injury. In Study 2, rmTBI once again increased alcohol consumption in female but not male rats, and repeated systemic treatment with JZL184 did not affect alcohol consumption. Also in Study 2, rmTBI increased anxiety-like behavior in males but not females and repeated systemic treatment with JZL184 produced an unexpected increase in anxiety-like behavior 6-8 days post-injury. In summary, rmTBI increased alcohol consumption in female rats, systemic JZL184 treatment did not alter alcohol consumption, and both rmTBI and sub-chronic systemic JZL184 treatment increased anxiety-like behavior 6-8 days post-injury in males but not females, highlighting robust sex differences in rmTBI effects.

## Introduction

Traumatic brain injury (TBI) is a heterogeneous disorder defined as an impairment in brain function resulting from an external force to the head. It is estimated that 69 million individuals will sustain a TBI worldwide each year, with 81% of all cases being of mild severity^1^. Repeated mild TBIs (rmTBI) and repeated sub-concussive head impacts are particularly relevant in those that play contact sports and in military veterans^2^. Repeated TBI leads to more severe symptoms and more prolonged recovery due to the cumulative effect of multiple injuries^3^. After TBI, cognitive dysfunction and psychiatric disorders are common and major contributors to long-term disability^4,5^. The prevalence of TBI tends to be higher in men than in women, but in certain subpopulations such as those participating in gender comparable sports, females have higher incidence of reported concussion than males^6–8^. There are sex differences in the quantity and severity of symptoms after repeated TBI. For example, one study reported that soccer players with a history of previous concussions performed worse than those who had not sustained a previous TBI, and also females soccer players performed worse on neurocognitive testing and reported more symptomatology compared to males^9^. Results from preclinical studies have also shown sex-dependent responses to repeated injury^10^.

Alcohol use disorder (AUD) is a major societal problem in the US, with more than 15 million AUD diagnoses in 2019^11^. TBI and alcohol use are highly comorbid, with studies showing up to 51% of TBIs are alcohol-related^12^. AUD is the third most common psychiatric disorder in people that have incurred a TBI^13^. Clinical studies suggest that alcohol consumption decreases in the first year after TBI^14,15^, but a subset of patients return to pre-injury levels by 2 year post-TBI^15^. Others report that patients with mild TBI return to alcohol use sooner post-injury than patients with moderate or severe TBI, possibly due to an earlier recovery of physical function^16^. It should also be acknowledged that many mild TBIs are not diagnosed^17^, and it is likely that individuals with undiagnosed mild TBIs consume alcohol acutely in the post-injury window.

Men have higher risk of post-TBI alcohol misuse compared to women^15^. Preclinical studies are mixed in terms of alcohol consumption patterns in male animals following TBI, with some studies showing that TBI increases consumption, sensitivity or motivation for alcohol^18–22^ while others report no increase or reduced alcohol intake after experimental TBI^20,22,23^. One preclinical study from our group tested operant alcohol self-administration in female rats after TBI and reported no effects of injury on that behavior^24^. A separate study showed that female mice (but not male mice) injured during adolescence self-administered more alcohol and showed increased alcohol Conditioned Place Preference (CPP) during adulthood compared to sham controls^22^. There is a need for studies examining sex differences in post-TBI alcohol drinking in preclinical models because there are sex differences in post-TBI alcohol use/misuse in humans ^25^, and because pre- and post-TBI alcohol use can worsen TBI outcomes (e.g., increased risk of seizures, neuropsychiatric disorders, and subsequent TBIs)^26–31^.

TBI and alcohol consumption each dysregulate endocannabinoid (eCB) function in the brain. eCBs are retrograde signaling molecules that exert biological effects via activation of CB1 and CB2 type receptors – the two major eCBs are 2-arachidonoyl glycerol (2-AG) and *N*-arachidonoylethanolamine (anandamide, AEA). These molecules are synthesized “on demand”, enhancing endocannabinoid tone in a site- and event-specific manner^32,33^. 2-AG is hydrolyzed mainly by monoacylglycerol lipase (MAGL, 85%) a serine-hydrolase enzyme mainly found in presynaptic terminals, and by α/β-hydrolase domain 6 and 12 (ABHD6, ABHD12, 15%), while AEA is hydrolyzed by fatty acid amide hydrolase (FAAH)^34^. Brain eCB dysregulation has been linked to various pathological conditions and therapeutic strategies, one of which is modulation of eCB signaling via manipulation of degradative enzymes such as MAGL and FAAH. JZL184 is a selective MAGL inhibitor and there is mounting evidence regarding the potential therapeutic effects of MAGL inhibition on post-TBI outcomes^18,35^. Other work has shown that treatment with MAGL inhibitors leads to attenuation of neurodegeneration, blood brain barrier (BBB) breakdown, lesion volume^33^, LTP impairments in the hippocampus^36^, and microglia activation^37^ after injury.

This work is timely and relevant due to the high number of individuals (with or without TBI) that use prescribed or unprescribed cannabinoid drugs to manage a wide array of symptoms such as pain, anxiety, and sleep disturbances^38^, and because of the interaction of cannabis with comorbid drugs such as alcohol.

The purpose of this study was to test the effects of repeated mild TBI on alcohol consumption and anxiety-like behavior, and to test the effects of repeated post-injury JZL184 treatment on post-TBI alcohol consumption and anxiety-like behavior in male and female rats.

## Methods

### Animals

A total of 112 Wistar rats, 4 months old at the time of TBIs (Charles Rivers Laboratories, Raleigh, NC, USA) were used (Study 1: males (n = 16) and females (n = 17); Study 2: males (n = 32) and females (n = 47)). Animals were housed under conditions of controlled temperature (70-71 F) and humidity (50%) and exposed to a 12-h reverse light-dark cycle. Standard rat chow food and water were provided *ad libitum*. Upon arrival, animals were allowed to acclimate to the colony room for one week. All procedures were approved by the Institutional Animal Care Use Committee of the Louisiana State University Health Sciences Center (LSUHSC; New Orleans, LA) and were conducted in accordance with the National Institute of Health (NIH) guidelines.

### TBI

The weight drop model consists of a U shape stage made with clear plastic (38 x 27 x 27 cm^3^) that contains a collection chamber containing a landing sponge. A clamp stand holds a plastic guide tube in place. TBIs is produced by dropping a weight of specified characteristics through a guide tube at a determined height onto the head of the animal^39^. Animals were anesthetized with isoflurane (4% induction and 2-3% maintenance). Bupivacaine (7mg/kg) was injected subcutaneously at the impact site to minimize pain during recovery. After being anesthetized, animals were placed in tin foil in a prone position, with their head directly beneath the bottom of the guide tube. A 400gr weight was released from 100cm height directly to the center of the head with the skin and skull remaining completely intact. Three TBIs were produced 24 hours apart. Immediately after each TBI righting reflex and respiratory rate were measured. Sham animals underwent an identical procedure, but the weight was not dropped. Animals were allowed to recover in their home cage. Animals were assigned to each group based on alcohol baseline.

### Drugs

Systemic injection of JZL184 results in MAGL inhibition and elevations of 2-AG in the brain up to 9-fold^40^. The maximal inhibition is achieved at 0.5 h post-treatment, persisting for at least 24 hours, with 85% inhibition of 2-AG hydrolysis activity^40^. In previous studies we showed that systemic JZL184 administration 30 min post TBI reduced neuroinflammation, improved neurobehavioral outcomes, and attenuated motivation for alcohol drinking in male rats^18,35,37^. JZL184 (Item 13158, Cayman Chemical, Ann Arbor, MI,) or vehicle was injected intraperitoneally (i.p.) 30 minutes after each TBI in two doses (16 mg/kg, after the first TBI, and 8 mg/kg, after the second and third TBI).

### Intermittent 2-Bottle Choice (I2BCH)

After arrival in the lab and an acclimatization period, animals were single-housed and provided access to ethanol (20% v/v) without sweeteners. Each rat was provided access to 1 bottle of 20% v/v ethanol and 1 bottle of water for 24 hours on Mondays, Wednesdays and Fridays followed by 24-48 hours with two water bottles on the home cage (Tuesdays, Thursdays, Saturdays and Sundays). The placement of the ethanol bottle was alternated in each ethanol drinking session to control for potential side preferences. Bottles were weighed before and after 24-hour access sessions. A bottle of water and alcohol placed in a cage without an animal was also weighed in each session to control for bottle leaks. Animals were weighed after each alcohol session to calculate the grams of ethanol intake per kilogram of body weight. Ethanol solutions were prepared from autoclaved water and 95% ethanol solution. After completing the 12 alcohol sessions pre-TBI, animals spent nine continuous days without alcohol, a time during which animals were subjected to Sham or TBI procedure, testing for NSS/NBS and recovery. The day ten after the last pre-TBI alcohol session, animals resumed alcohol consumption for 12 more sessions.

### Neurological Behavioral Score

Twenty-four hours after the last TBI (Fig. 1) we tested neurological and neurobehavioral deficits using Neurological Severity Score (NSS) and Neurobehavioral Score (NBS, Fig. 1). NSS evaluates somatomotor and somatosensory functions through a series of tasks involving motor, sensory reflexes, balancing, and motor coordination, with total scores ranging from 0 to 25. NBS was tested immediately after NSS; this test measures exploratory and motor behavior, lateral pulsion, and object recognition, (see^41^), with total scores ranging from 0 to 12. Higher scores indicate more performance deficits in both tests.

**Figure 1.**
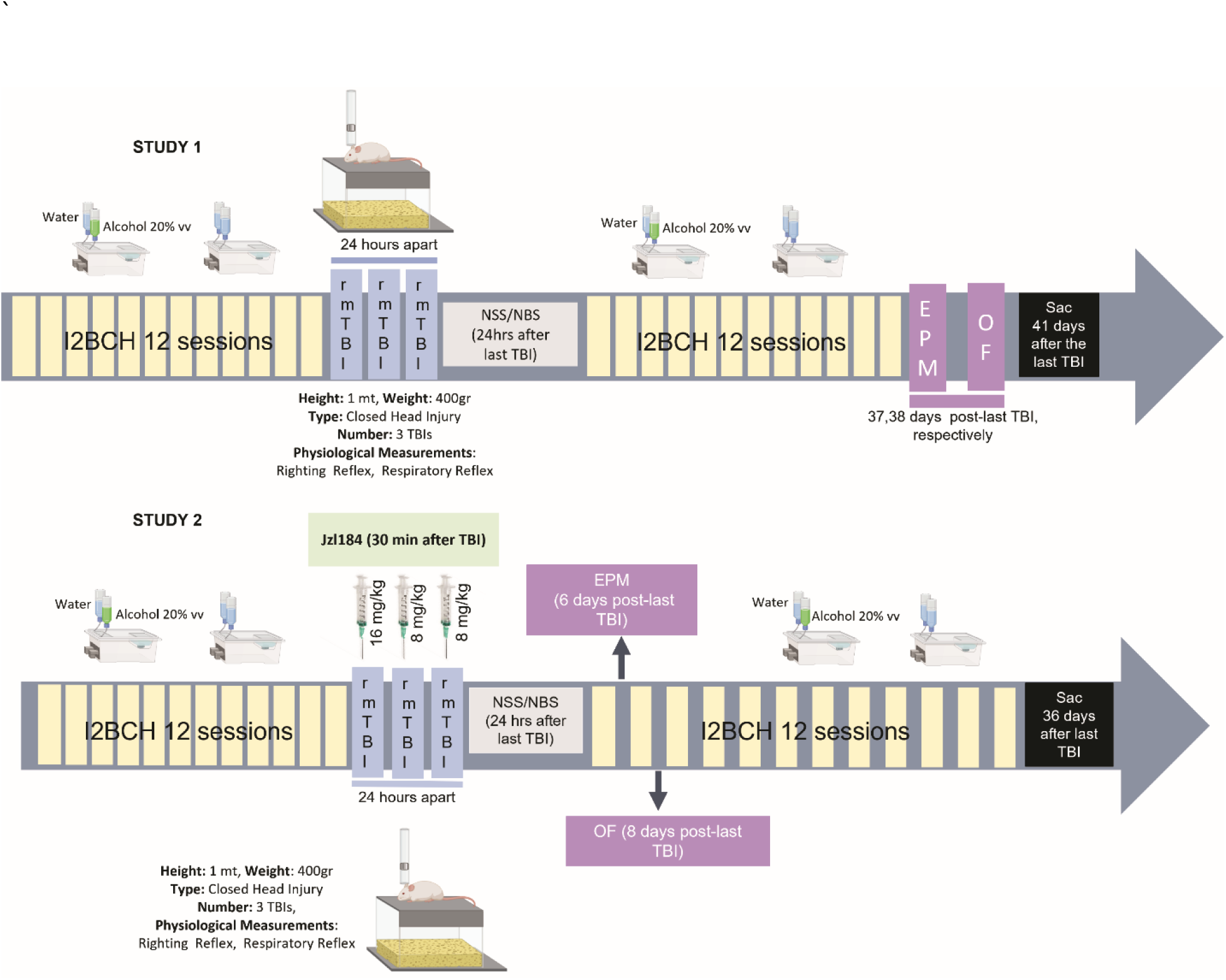
Timeline for Study 1 and 2. In Study 1, male and female adult Wistar rats were exposed to alcohol consumption using I2BCH. Two bottles, one containing 20% v/v alcohol and one containing water were placed in the cages three days per week (Mondays, Wednesdays, and Fridays) while two bottles of water were placed on Tuesdays, Thursdays, and weekends. Animals were exposed to alcohol for a total of 12 sessions pre- and 12 sessions post-rmTBI. Immediately after each TBI, respiratory rate and righting reflex were measured. Twenty-four hours after the last TBI animals were tested on NSS and NBS. Anxiety-like behavior was tested using EPM and OF on days 37 and 38 post-last TBI. In Study 2, male and female adult rats were exposed to the same parameter of alcohol consumption than Study 1. Thirty minutes after each TBI animals were injected with JZL184 (16, 8, and 8 mg/kg). Immediately after each TBI, respiratory rate and righting reflex were measured. Twenty-four hours after the last TBI animals were tested on NSS/NBS. Six and eight days after the last TBI animals were tested for anxiety-like behavior using EPM and OF. Portions of this figure were created with biorender.com

### Blood Alcohol Levels (BAL)

Blood samples were collected by tail snip 30 minutes after the commencement of the last alcohol session. The tip of the tail (∼1 mm) was cut with a disposable scalpel, and blood was collected into a microcentrifuge tube. Topical bupivacaine was used to avoid pain after bleeding. BAL was analyzed using an ANALOX machine according to the manufacturer’s instructions (Analox Instruments Limited, London, England).

### Elevated Plus Maze (EPM)

In Study 1, animals were tested for anxiety-like behavior in the elevated plus-maze (EPM) 3 days after the last alcohol session, approximately 37 days after the last TBI, and 3-4 h into the dark cycle. In Study 2, animals were tested in the EPM 6 days after the last TBI, and testing occurred 8 hours into the dark cycle, on a day when the animals did not consume alcohol. Rats were individually placed in the center of the EPM facing one of the open arms and allowed 5 min to explore the maze. Sessions were recorded by a video camera placed directly above the apparatus. The apparatus was cleaned with quatricide between animals to avoid olfactory cues. The percentage of total time spent in the open arms was calculated as [open arms time (sec)/ open arms time + closed arms time (sec)] x 100. Open arm entries (another index of anxiety-like behavior), closed arm entries, and total arms entries were recorded when all four paws were placed in the arm. An arm exit was recorded when at least two paws were out of the arm.

### Open Field (OF)

The open field consists of a square-shaped methacrylate chamber (78 x 78cm) with the bottom divided into 25 squares (15 x 15cm). In Study 1, animals were tested 38 days after the last TBI, and 3-4 hours into the dark cycle. In Study 2, animals were tested 8 days after the last TBI, and 8 hours into the dark cycle on a day when animals did not consume alcohol. Animals were placed in the arena for 5 min, and a ceiling-mounted video camera recorded the session. The amount of time each rat spent exploring the center (defined by exploring boxes that do not border the walls of the apparatus) and periphery (defined by exploring boxes that border the walls of the apparatus) was quantified. The percentage time spent in the center was calculated as [(seconds in the center /300) X 100], and lower scores in this measure were interpreted as higher anxiety-like behavior. The number of lines crossed was calculated and interpreted to reflect non-specific locomotor activity. The apparatus was cleaned with quatricide between animals.

### Statistical Analysis

Statistical analyses were performed with SPSS (IBM Corp., Armonk, NY, USA). Outliers were identified using quartiles (Q) and interquartile range (IQR) and those animals with values lower than Q1-1.5*IQR or higher than Q3+1.5*IQR were excluded from the corresponding analysis. One male from the TBI vehicle-treated, and one male from the Sham JZL184-treated group were removed from the analysis of EPM. One female from the Sham vehicle-treated group was removed from the analysis of the OF. In Study 1, respiratory rate and righting reflex were analyzed with 3-way repeated measure ANOVAs (RM ANOVAs) (between-subject factors: injury and sex; within-subject factor: time). NSS and NBS data were analyzed with 2-way ANOVAs (between-subjects factors: injury and sex). Alcohol consumption per day and total alcohol consumption per week were analyzed using 3-way RM ANOVAs (between-subject factors: injury and sex; within-subject factor: time). Cumulative alcohol consumption, BAL, EPM, and OF were analyzed with two-way ANOVAs (between-subjects factors: injury and sex). In Study 2, respiratory rate and righting reflex were analyzed with 3-way RM ANOVAs (between-subject factors: injury and sex; within-subject factor: time). NSS/NBS, EPM, OF, and cumulative alcohol consumption were analyzed with three-way ANOVAs (between-subjects factors: injury, sex, JZL184 treatment). Alcohol consumption per day and alcohol consumption per week were analyzed with 3-way RM ANOVAs (between-subject factors: injury and JZL184; within-subject factor: time). *Post hoc* comparison between pairs of groups was performed with Sidak when indicated by omnibus ANOVA results. Statistical significance was set as p = 0.05. All data are expressed as mean ± standard error of the mean (SEM).

## Results

### Repeated mild TBI does not affect physiological response

Immediately after each TBI, respiratory rate and righting reflex were measured to determine the effect of rmTBI on the physiological parameters. In Study 1, a 3-way RM ANOVA revealed that females exhibited lower respiratory rate than males (F_(1,29)_ = 31.61, p = <0.001), but there were no other main effects or interactions effects (Fig. 2A). A separate 3-way RM ANOVA revealed a main effect of time on righting reflex such that time to recover righting reflex decreased over time (F_(2,58)_ = 4.77, p = 0.012), but there were no other main effects or interactions effects (Fig. 2B). In Study 2, a 3-way RM ANOVA also revealed lower respiratory rate in females relative to males (F_(1,75)_ = 42.57, p = p = < 0.001), but there were no other main effects or interactions effects (Fig. 2C). A separate 3-way RM ANOVA revealed a time x injury interaction effect (F_(2,150)_ = 3.86, p = 0.023) on righting reflex such that in the third TBI, the Sham Control group is significant different from the TBI group and, and there are significant differences between the first and the third TBI in the Sham control group (Fig. 2D). There were no other main effects or interactions effects on righting reflex in Study 2.

**Figure 2.**
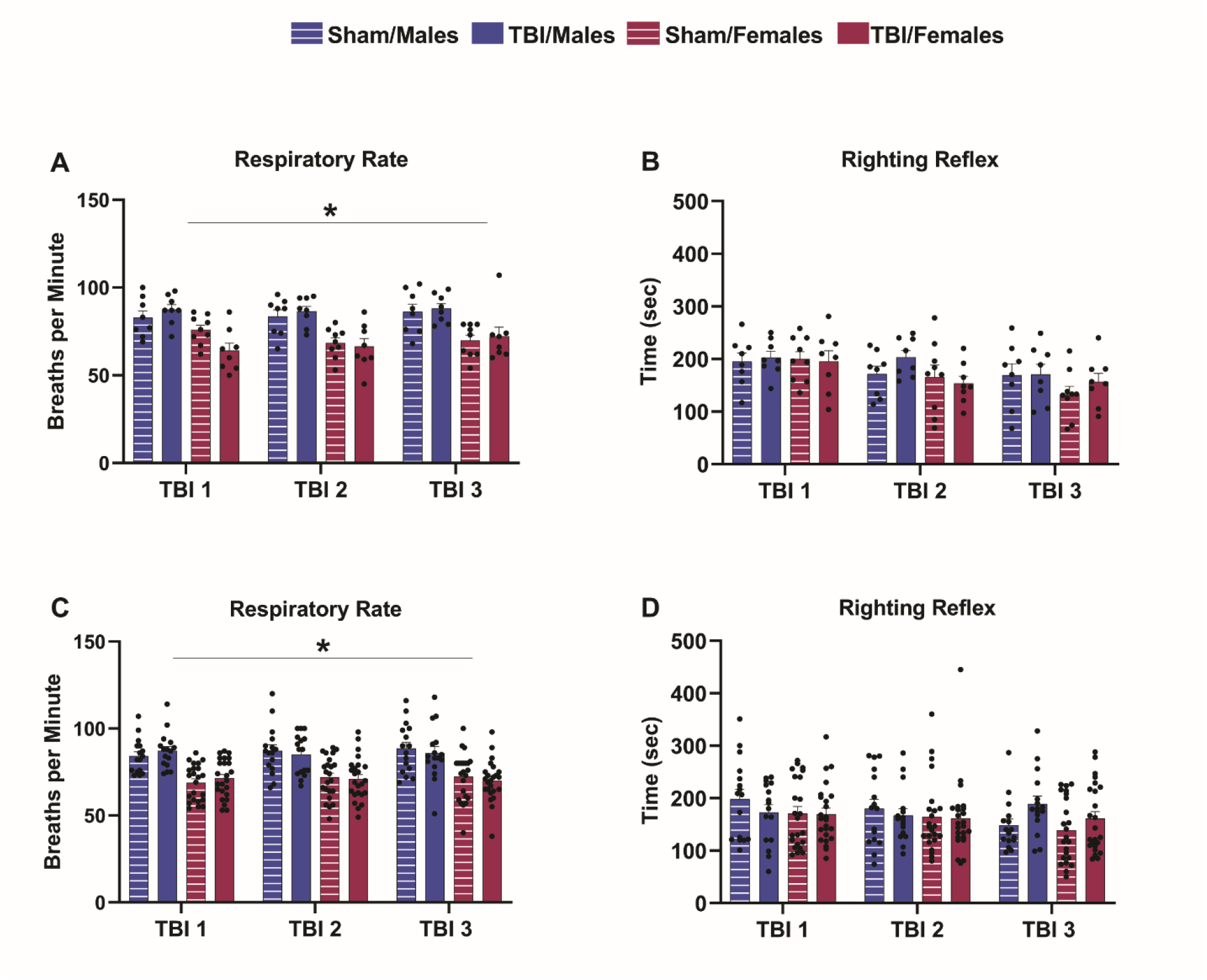
Repeated mild TBI does not affect physiological response. Respiratory rate (breaths per minute) (A, B) and righting reflex (time to recover the loss of consciousness in seconds) (C, D) were tested immediately after each TBI. Data were analyzed via RM ANOVA. (*) for males compared to females. Study 1, n = 8-9 per group; Study 2, 8-12 per group.

### Repeated mild TBI increases NSS and NBS Scores

To determine the effects of rmTBI on NSS and NBS, we tested animals 24 hours after the last TBI. In Study 1, a 2-way ANOVA revealed higher NSS scores in TBI rats relative to Sham Controls (F_(1,29)_ = 11.05, p = 0.002), but there was no effect of sex nor an injury x sex interaction effect (Fig. 3A). A separate 2-way ANOVA revealed no main effects or interaction effects of injury and sex on NBS scores (Fig. 3B). In Study 2, a 3-way ANOVA again revealed higher NSS scores in TBI rats relative to Sham Controls (F_(1,71)_ = 28.75, p = <0.001), and males also exhibited higher NSS scores than females (F_(1,71)_ = 4.61, p = 0.035). There were no other main effects or interaction effects of any factor on NSS scores in Study 2 (Fig. 3C). A separate 3-way ANOVA revealed higher NBS scores in TBI rats relative to Sham Controls (F_(1,71)_ = 6.96, p = 0.010), but no other main effects or interaction effects (Fig. 3D).

**Figure 3.**
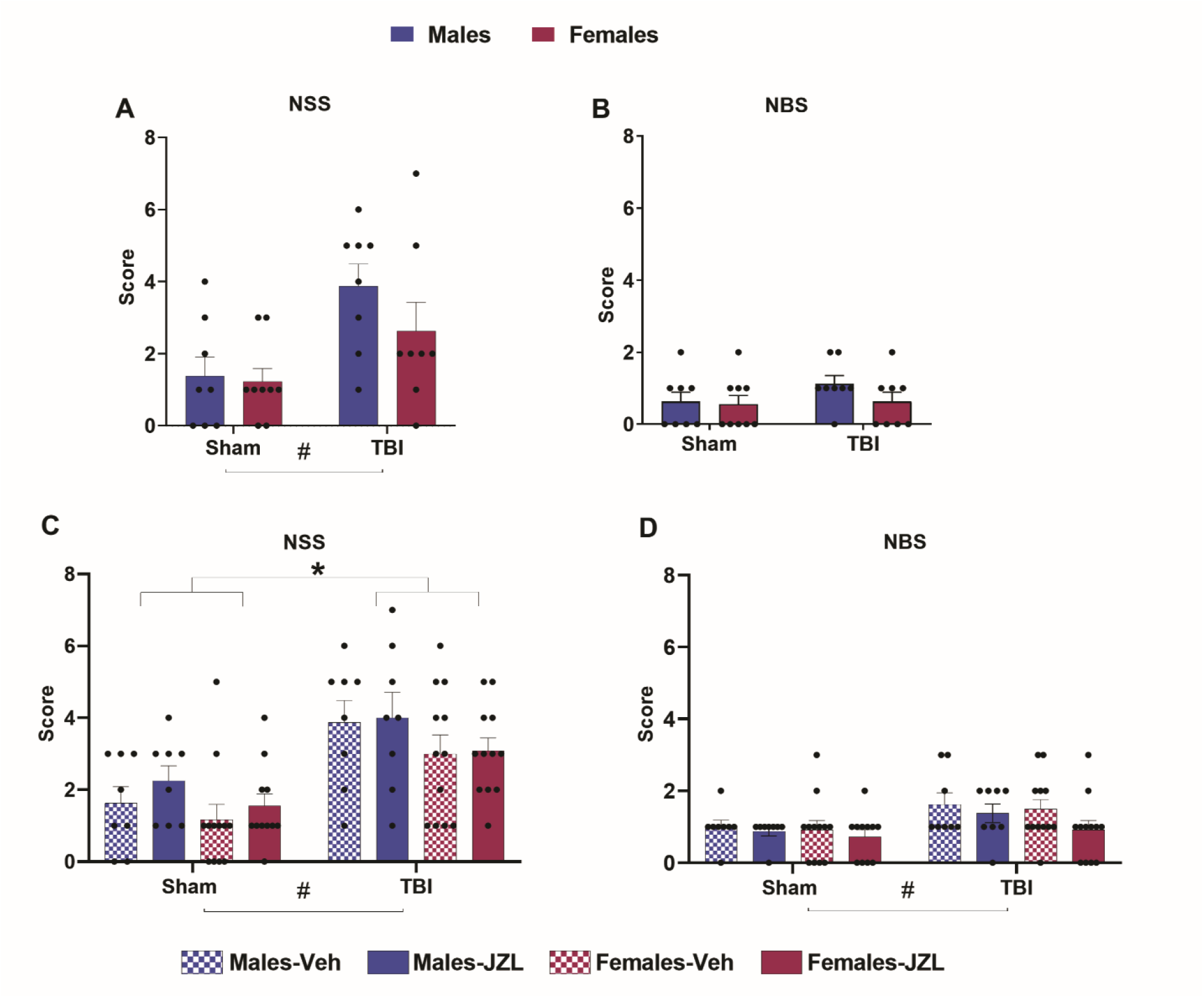
Repeated mild TBI increases NSS and NBS Scores. In Study 1, Neurological Severity and Behavioral Scores were tested 24 hours after the last Sham or TBI procedures (A, B). In Study 2, animals were treated with 3 systemic injections of JZL184, 30 minutes after each TBI, and tested on the NSS/NBS 24 hours after the last Sham or TBI procedures (C, D). Data were analyzed with 2- and 3-way ANOVA, respectively. (#) for Sham compared to TBI, (*) for males compared to females. Study 1, n = 8-9 per group; Study 2, n = 8-12 per group.

### Repeated mild TBI increases alcohol consumption in females

Animals were allowed to drink alcohol using an I2BCH procedure (described above and in Figure 1) to test rmTBI effects on alcohol consumption in males and females. Alcohol consumption pre-TBI in males and females is shown in Fig. 4A. When alcohol post-TBI is analyzed, a 3-way RM ANOVA of daily drinking data revealed a main effect of sex (F_(1,29)_ = 18.09, p = <0.001) and an injury x sex interaction effect, (F_(1,29)_ = 12.80, p = 0.001), such that TBI females consumed more alcohol than Sham females and TBI males (p < 0.05 in both cases). There was a main effect of time on daily alcohol consumption (F_(12,348)_ = 3.12 p = <0.001) (Fig. 4B). There were no other main effects or interaction effects on daily alcohol drinking in Study 1. When analyzed by total consumption per week, a 3-way RM ANOVA revealed significantly higher alcohol consumption in females relative to males (F_(1,29)_ = 18.07, p = <0.001), and an injury x sex interaction effect (F_(1,29)_ = 13.94, p = <0.001), such that TBI females consumed more alcohol than Sham females and TBI males (p < 0.05 in both cases) (Fig. 4C). There was also a main effect of time (F_(3,87)_ = 3.25, p = 0.025) such that animals consumed less alcohol over time. There were no other main effects or interaction effects on weekly alcohol drinking in Study 1. Finally, we analyzed cumulative alcohol consumption during the 12 post-TBI I2BCH sessions, and a 2-way ANOVA revealed once again that females consumed more alcohol than males (F_(1,29)_ = 18.07, p = <0.001), as well as an injury x sex interaction effect (F(_1,29)_ = 13.94, p = 0.001), such that TBI females consumed more alcohol than Sham females and TBI males (p < 0.05 in both cases) (Fig. 4D). There were no other main effects or interaction effects on cumulative post-TBI alcohol drinking in Study 1. We measured BALs 30 minutes after the commencement of the last alcohol session and 2 -way ANOVA revealed higher BALs in TBI animals (F_(1,29)_ = 5.70, p = 0.024), and an injury x sex interaction effect (F_(1,29)_ = 4.52, p = 0.042) such that TBI females exhibited higher BALs than Sham females and TBI males (p < 0.05 in both cases) (Fig. 4E).

**Figure 4.**
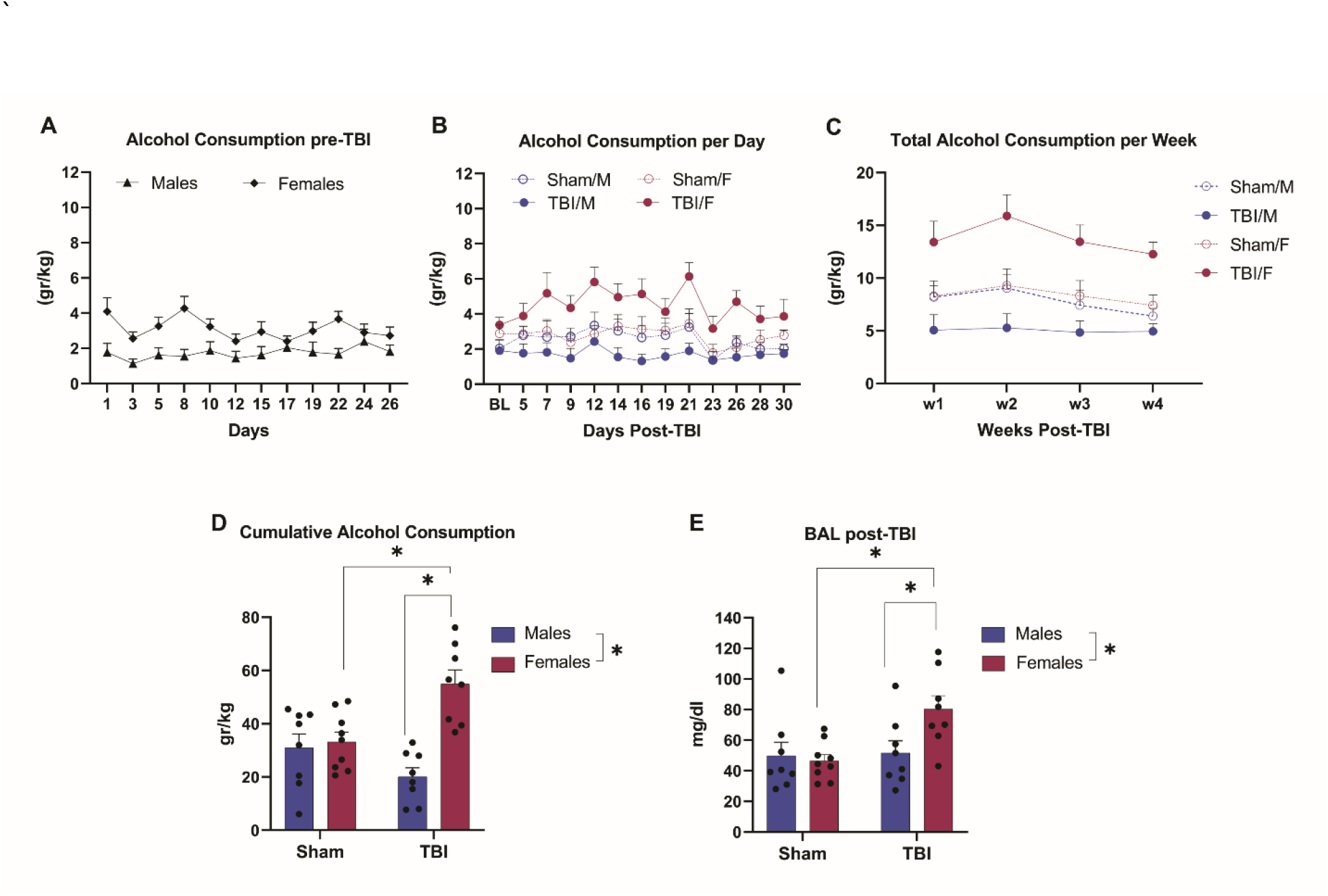
Repeated mild TBI increases alcohol consumption in females. Alcohol consumption pre-rmTBI, per day (gr/kg) in males and females, 12 sessions using I2BCH (A). Alcohol consumption post-rmTBI, per day (gr/kg) in males and females subjected to Sham or rmTBI, 12 sessions (B). Total alcohol consumption per week (gr/kg), over 4 weeks, in males and females subjected to Sham or rmTBI (C). Cumulative alcohol consumption (gr/kg) over 12 sessions post-TBI, in males and females subjected to Sham or rmTBI (D). Blood alcohol levels (mg/dl) were tested 30 minutes after the commencement of the last alcohol session for the Sham and rmTBI groups (E). n = 8-9 per group.

### rmTBI does not increase long-term anxiety-like behavior

To determine whether rmTBI increases long-term anxiety-like behavior, we tested animals in the EPM 37 days post-last TBI. A 2-way ANOVA revealed a main effect of sex on the percentage of time exploring the open arms of the EPM (F_(1,29)_ = 25.91, p = <0.001). There were no other main effects or interaction effects (Fig. 5A). Analysis of the number of entries in the open arms by 2-way ANOVA revealed a main effect of sex (F_(1,27)_ = 17.43, p = < 0.001) on the number of entries in the open arms of EPM. There were no other main or interaction effects (Fig. 5B). A 2-way ANOVA revealed a main effect of sex (F_(1,27)_ = 5.03, p = 0.033) on the number of entries in the closed arms. There were no other main or interaction effects (Fig. 5C). When total arms entries were analyzed, a 2-way ANOVA showed a main effect of sex (F_(1,27)_ = 14.36, p = 0.001). There were no other main or interaction effects (Fig. 5D).

**Figure 5.**
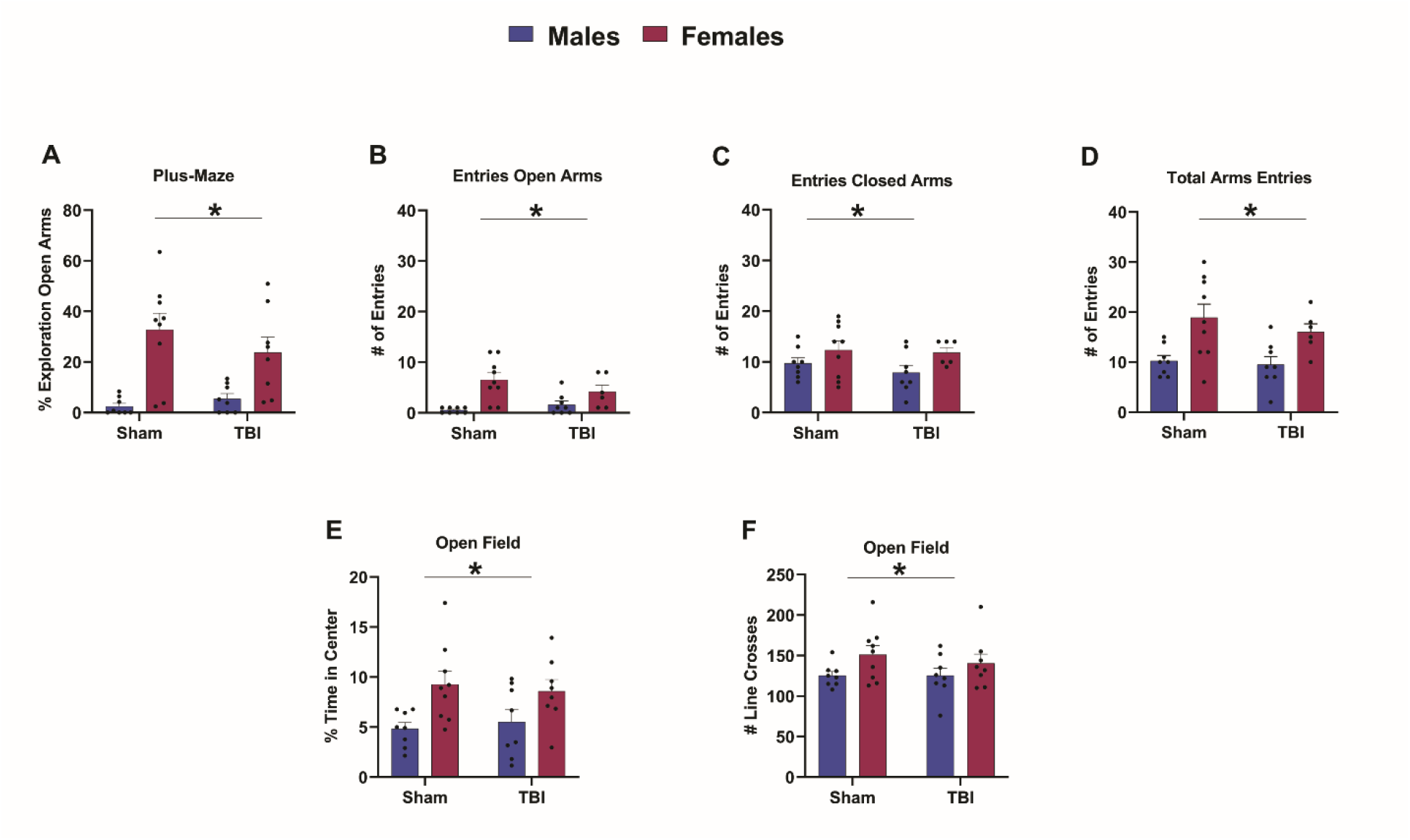
Repeated mild TBI does not alter long-term anxiety-like behavior. Anxiety-like behavior in the elevated plus maze and the open field 37 and 38 days post-TBI. Percentage of time exploring the open arms in the EPM, 37 days post-rmTBI, for males and females subjected to Sham or rmTBI (A). The number of entries in the open (B), closed arms (C) and total arm entries (D) are shown. Percentage of time in the center of the open field 38 days post-TBI (E) and the number of lines crossed for male and female rats subjected to Sham or rmTBI (F). Data were analyzed with 2-way ANOVA. (*) for males compared to females. n = 8-9 per group.

Animals were tested in the OF at 38 days post-last TBI. 2-way ANOVA showed a main effect of sex on the percentage of time spent in the center of the OF (F_(1,29)_ = 10.91, p = 0.003). There were no other main effects or interaction effects on the percentage of time spent in the center of the OF (Fig. 5E). A 2-way ANOVA showed a main effect of sex (F_(1,29)_ = 4.54, p = 0.042) in the number of lines crossed. There were no other main effects or interaction effects (Fig. 5F).

### Repeated systemic JZL184 treatment induces short-term anxiety-like behavior

In Study 2, to determine the effect of rmTBI and repeated systemic JZL184 treatment on anxiety-like behavior, male and female rats were tested on the EPM 6 days after the last TBI. Each of the three systemic JZL184 treatments occurred acutely post-injury on the three TBI days. A 3-way ANOVA showed a main effect of TBI to reduce time exploring the open arms of the EPM (F_(1,69)_ = 6.02, p = 0.017), as well as a surprising main effect of JZL184 to reduce time exploring the open arms of the EPM (F_(1,69)_ = 5.19, p = 0.026), and finally a main effect of sex such that males spent less time in the open arms of the EPM (F_(1,69)_ = 12.82, p = 0.001). There were no interaction effects on time spent in open arms of the EPM (Fig. 6A). A separate 3-way ANOVA revealed that TBI animals entered the open arms of the EPM fewer times than Sham Controls (F_(1,69)_ = 5.85, p = 0.018), and male rats entered the open arms of the EPM fewer times than female rats (F_(1,69)_ = 16.46, p = < 0.001). There were no other main effects or interaction effects on open arm entries (Fig. 6B). A separate 3-way ANOVA revealed an injury x sex interaction effect (F_(1,69)_ = 6.76, p = 0.011) and an injury x JZL x sex interaction effect (F_(1,69)_ = 4.52, p = 0.037) on closed arm entries in the EPM. Post-hoc analyses revealed that number of entries by Sham Control males is different from Sham Control females. In the JZL184-treated group, number of entries by Sham Control males is different from TBI males, and different in Sham Control females from TBI females. In the TBI group, number of entries by the JZL184-treated males differed from JZL184-treated females. In the Sham Control group, number of entries by JZL184-treated males differed from JZL184-treated females (Fig. 6C). When total entries to the open and closed arms were analyzed, a 3-way ANOVA showed a main effect of sex (F_(1,69)_ = 12.31, p = 0.001). There were no other main effects. There was also an injury x sex (F_(1,69)_ = 4.28, p = 0.042) and an injury x JZL x sex interaction effect (F_(1,69)_ =6.13, p = 0.016) (Fig. 6D). Post-hoc analyses revealed the total arms entries by Sham Control females is different from TBI females, and that Sham Control males are different from Sham Control females. Post-hoc analysis also showed that in the JZL184-treated group Sham Control females are different from TBI females, in the Sham Control group, JZL184-treated females are different from JZL184-treated males, and in the TBI group, vehicle-treated females are different from vehicle-treated males.

**Figure 6.**
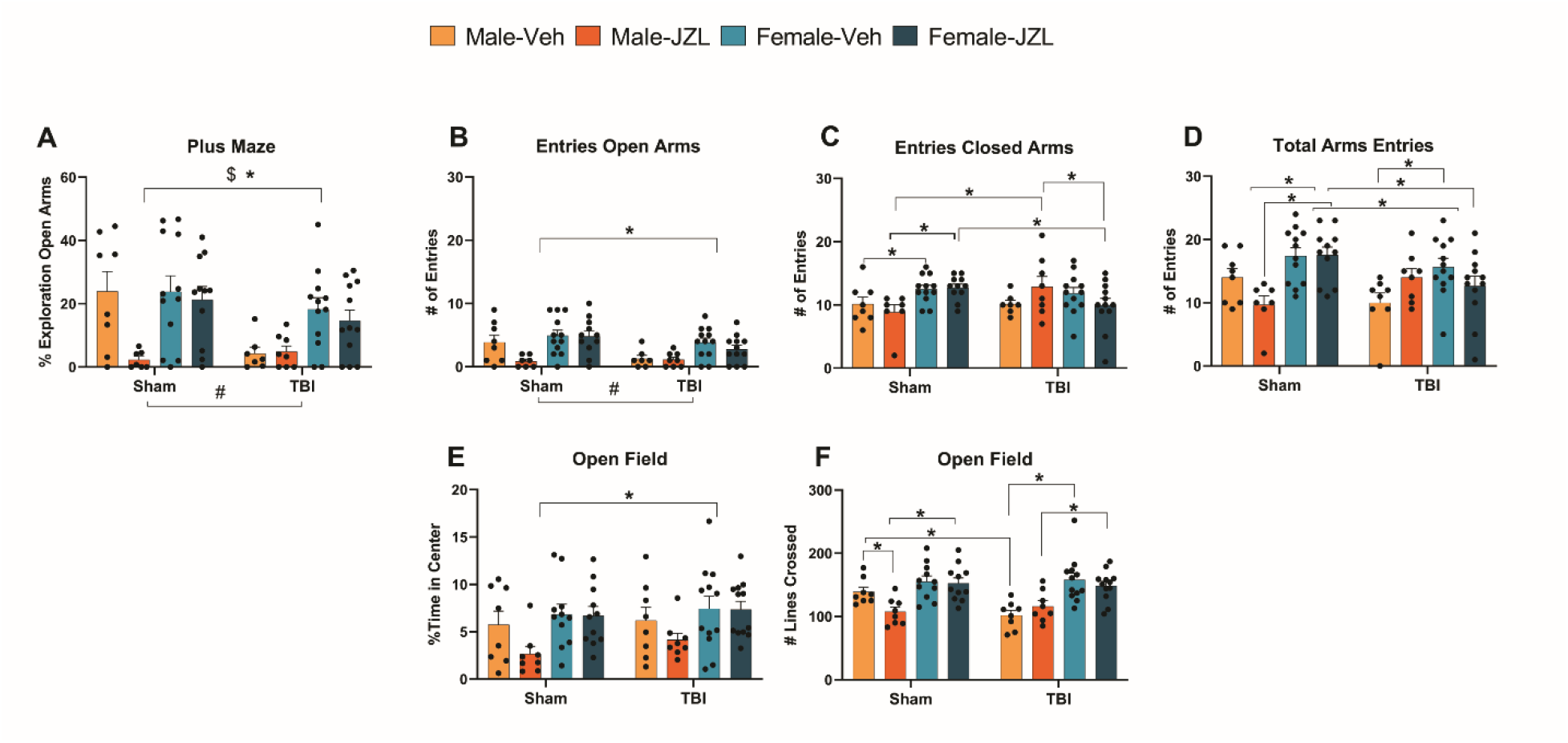
JZL184 induces short-term anxiety-like behavior. Anxiety-like behavior in the elevated plus maze and the open field 6 and 8 days post-TBI. Percentage of time exploring the open arms in the EPM 6 days post-rmTBI, for males and females subjected to Sham or rmTBI and treated with 3 systemic injections of JZL184, 30 minutes after each TBI (A) Number of entries in the open (B), and closed arms (C), and total arms entries (D) are shown. Percentage of time in the center of the open field 8 days post-TBI in males and females subjected to Sham or rmTBI and treated with 3 systemic injections of JZL184, 30 minutes after each TBI (E) and a number of lines crossed (F). Data were analyzed with 3-way ANOVA. For percentage exploration in open arms and percentage time in the center: (*) denote comparison between males and females, (#) denote comparison between Sham control and TBI and ($) denote comparison between vehicle and JZL184. For number of entries in the open arm, closed arm, and total arms entries, and number of lines crossed (*) denote differences between the specific groups. n = 8-12 per group.

In the Open Field test, a 3-way ANOVA revealed that male rats spent less time in the center of the OF than female rats (F_(1,70)_ = 9.00, p = 0.004), and there were no other main effects or interaction effects on this outcome measure (Fig. 6E). A separate 3-way ANOVA showed male rats crossed fewer lines than female rats during the 5-minute test (F_(1,70)_ = 36.73, p = < 0.001). There was an injury x JZL x sex interaction effect on the number of lines crossed (F_(1,70)_ = 4.47, p = 0.038). Post hoc analyses showed that in the vehicle groups, number of line crosses in Sham Control males differed from TBI males. In the Sham Control group, number of line crosses in vehicle-treated males were significantly different from JZL184-treated males, and JZL184-treated males and different from JZL184-treated females. In the TBI group, number of line crosses by JZL184-treated males differed from JZL184-treated females and between vehicle-treated males and vehicle-treated females. There were no other main effects or interaction effects on this outcome measure (Fig. 6F).

### Repeated systemic JZL184 treatment does not reduce alcohol consumption in females

In Study 2, we tested the effect of repeated systemic JZL184 treatment (acutely post-injury on the three mTBI days) on post-TBI alcohol consumption using the I2BCH procedure. Alcohol consumption pre-TBI in males is shown in Fig. 7A. A 3-way RM ANOVA revealed a main effect of time on alcohol consumption (F_(12,336)_ = 3.54, p = <0.001), but no other main effects or interaction effects on this outcome measure (Fig. 7B). When we analyzed weekly alcohol consumption data, we again observed a main effect of time (F_(3,84)_= 6.56, p = < 0.001), but no other main effects or interaction effects (Fig. 7C). When we used a 2-way ANOVA to analyze cumulative alcohol consumption during the 12 post-TBI sessions, we observed no effect of TBI or JZL184 treatment, nor an interaction effect (Fig. 7D).

**Figure 7.**
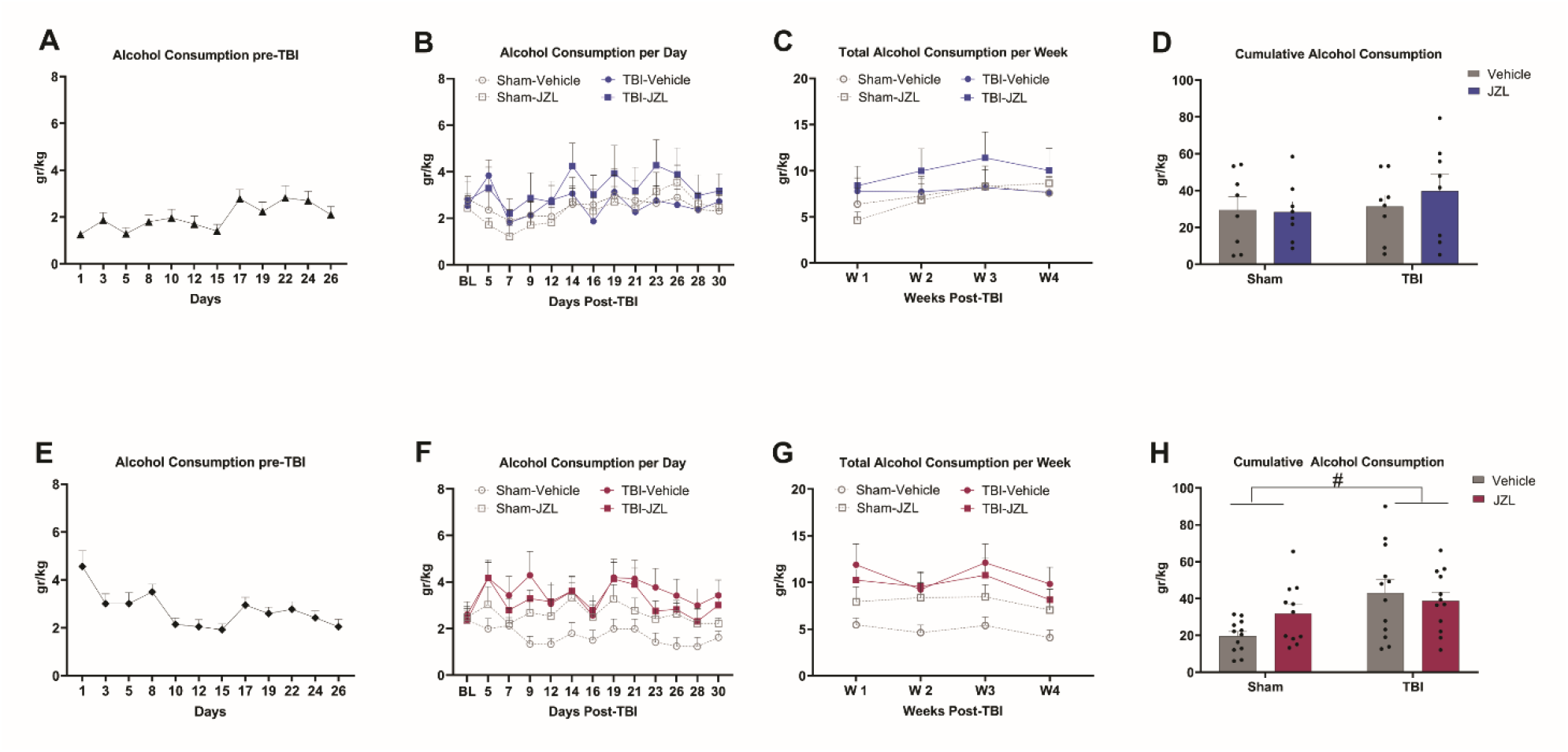
JZL184 does not reduce alcohol consumption in females. Alcohol consumption pre-rmTBI, per day (gr/kg) in males (A) and females (E) using I2BCH. Alcohol consumption post-rmTBI, per day (gr/kg) in males (B) and females (F) subjected to Sham or rmTBI and treated with 3 systemic injections of JZL184, 30 minutes after each TBI. Total alcohol consumption per week (gr/kg), over 4 weeks, males (C) and females G. Cumulative alcohol consumption (gr/kg) over 12 sessions post-TBI, males (D) and females (H). Data were analyzed with RM ANOVA. (#) for Sham compared to TBI. n = 8-12 per group.

Alcohol consumption pre-TBI in females is shown in Fig 7E. A 3-way RM ANOVA revealed a main effect of time on alcohol consumption (F_(12,516)_ = 3.14, p = < 0.001), and a main effect of TBI to increase alcohol consumption in females (F_(1,43)_ = 8.09, p = 0.007), but no other main effects or interaction effects on this outcome measure (Fig. 7F). When we analyzed weekly alcohol consumption data, we again observed a main effect of time (F_(3,129)_ = 4.44, p = 0.005) and higher alcohol consumption in TBI females (F_(1,43)_ = 8.68, p = 0.005), but no other main effects or interaction effects (Fig. 7G). When we used a 2-way ANOVA to analyze cumulative alcohol consumption during the 12 post-TBI sessions, we again observed higher alcohol consumption in TBI females (F_(1,43)_ = 8.69, p = 0.005), but no main effect of JZL184 treatment and no interaction effect (Fig. 7H).

## DISCUSSION

The purpose of the present study was to test the effect of rmTBI (using a weight drop model) on alcohol consumption and anxiety-like behavior in male and female rats (Study 1), and to determine the role of repeated systemic JZL184 treatment on post-TBI alcohol consumption and anxiety-like behavior in male and female rats (Study 2). We spaced repeated injuries by 24 hours because injuries that occur close together in time produce greater cognitive, histological, and behavioral impairment than injuries separated by longer periods of time and single injuries^42–48^.

In Study 1 and Study 2, we did not observe a significant alteration in physiological responses following rmTBI. One of the clinical signs to classify the severity of TBI is the duration of loss of consciousness. Mild TBI is characterized by no or a transient loss of consciousness (0-30 minutes)^49^. The righting reflex is a measure used as a surrogate of loss of consciousness in humans and is an indicator of injury severity and a righting reflex under 10-15 min has been suggested to reflect mild TBI severity^50^. The righting reflex in our studies was under 10 minutes, consistent with a mild TBI. We did observe sex differences in respiratory rate (but no effect of rmTBI), with females having fewer breaths per minute compared to males, which may suggest increased sensitivity to the effect of anesthesia in females compared to males.

When we tested animals in NSS and NBS assays 24 hours after the last injury, we found neurological deficits in animals subjected to rmTBI. In Study 2, animals were treated with systemic JZL184 or vehicle, 30 min after each Sham or mTBI procedure, and the drug did not affect post-TBI neurological outcomes. This is contrast to our previous findings in which systemic JZL184 treatment 30 min after lateral fluid percussion injury reduced post-TBI neurological deficits^35,37^. This discrepancy could be attributed to the different model used for TBI in those studies (LFP) or the type of TBI (single vs repeated). In the LFP a direct injury is inflicted in the brain by the application of a fluid pressure pulse^51^. By inducing a single TBI, we found that JZL184 attenuated neurological deficits up to 2 weeks post-injury^35^. However, whether we would find the same effects after repeated TBI by LFP remains unknown. The weight drop is a closed head injury in which the lesion is caused by a weight that is released from a height hitting the head of the animal. Animals were subjected to 3 TBIs and it is known that repeated TBI has cumulative effect on the brain and is related with more persistent symptomatology^52,53^. It is possible that the neurological deficits after single LFP or even after a single WDM, are less resistant to treatment compared with those produced by repeated TBI.

The I2BCH is a voluntary model of alcohol consumption that provides the animal with the opportunity to choose when and how much alcohol to drink^54^. Some studies have shown that rats consume substantial amounts (5-6gr/kg/24h) of 20% v/v alcohol, with approximately 30% of animals reaching high BAL (90-100 mg/dL)^55^. The I2BCH model has been used to mimic binge-like alcohol drinking in rodents^54^.

Most preclinical studies on post-TBI alcohol consumption have used male animals and have yielded mixed results, with some showing that experimental TBI does not increase alcohol consumption^20,23,24^, and others showing increased alcohol consumption, preference, or motivation following TBI^18,19,21^. Similarly, prior work using repeated TBI in animal models have reported increases in “front-loading” of alcohol consumption^56^ or no change in alcohol consumption^57^.

Sex differences in outcomes after TBI have been reported in humans and rodents (reviewed by^58,59^), but studies on sex differences in alcohol consumption after TBI are sparse. Here, in Study 1, we showed that rmTBI increases two-bottle choice alcohol consumption in females but not males, and blood-alcohol levels were accordingly higher in female rmTBI rats. Prior work from our group reported that a single lateral fluid percussion TBI increases operant alcohol self-administration in adult male Wistar rats^21^, but not in adult female Wistar rats^24^.

In Study 1, we did not detect rmTBI effects on anxiety-like behavior 37-38 days later in males or females. Prior work in male mice has reported increases in anxiety-like behavior at one year after repeated TBI (30 impacts, once per day, 5 days per week) induced by a diffuse TBI model^60^. One study in male mice reported increases in anxiety-like behavior 14 days but not 1-6 months after rmTBI (42 impacts, 6 per day, each hit separated by 2 h) induced by a controlled cortical impact^61^. It is possible that our test time point in Study 1 was too late to detect TBI-induced increases in anxiety-like behavior. In Study 2, we observed rmTBI-induced increases in anxiety-like behavior in the elevated plus-maze (tested 6 days after the last TBI), but not in the open field test (tested 8 days after the last TBI). Similar to Study 1, we observed higher anxiety-like behavior in males compared to females. All rats in our studies consumed alcohol for 12 sessions pre-rmTBI and 12 sessions post-rmTBI. One study reported that young adult alcohol-drinking (pre-TBI, or pre- and post-TBI consumption) female rats exhibited less post-TBI anxiety-like behavior in the elevated plus-maze compared to alcohol-naïve females^62^. Another study reported that male mice subjected to weight drop TBI, and treated with a high dose of alcohol (5g/kg) 30 minutes before injury exhibited less anxiety-like behavior in the open field test relative to saline-injected mice^63^. In our study, females consumed significantly more alcohol than males and exhibited less anxiety-like behavior than males, which may be interpreted as a protective effect of alcohol. Another consideration in our data is that animals were single-housed to measure home cage alcohol consumption, therefore it is possible that males were more susceptible to the effect of this housing condition on anxiety-like behaviors, although prior work reported that social isolation is more stressful for females while crowding is more stressful for males^64^.

Considerable data exists on the involvement of eCBs in regulating anxiety-like behavior in rodents. For example, acute pharmacologic inhibition of the degradative enzymes FAAH or MAGL reduces anxiety-like behavior both in rats and in mice^65–67^, especially under aversive conditions^68,69^. Conversely, genetic deletion or pharmacological blockade of CB1 receptors increases anxiety-like behavior^70–72^. Previous studies in our lab reported that systemic treatment with JZL184 (16 mg/kg) 30 minutes after a single lateral fluid percussion TBI reduces anxiety-like behavior in the open field test 7 days later^18^.

In Study 2, we systemically treated animals with JZL184 thirty minutes after each of three rmTBI injuries, and we tested rats for alcohol consumption over days after the last injury/treatment combination. We did not observe systemic JZL184 effects on post-TBI two-bottle choice alcohol consumption. Prior studies reported that synthetic cannabinoid receptor agonists increase alcohol consumption in Sardinian alcohol preferring rats^73^, increases the motivation for alcohol consumption^74^ and increases alcohol seeking and alcohol self-administration during relapse^75^ and increases relapse-like alcohol drinking in rats^76^. Conversely, systemic treatment with the cannabinoid receptor antagonist reduces operant alcohol responding and two bottle choice drinking in outbred rats^77–79^ and inbred mice^80^, reduces operant alcohol responding, two-bottle choice alcohol drinking, and relapse like alcohol-seeking behavior in rats selectively bred for high alcohol preference^81–83^, reduces operant alcohol self-administration and free-choice drinking in alcohol dependent rats^84,85^, prevents footshock stress-induced increases in alcohol preference in mice^86^, and CB1 receptor knockout mice experience less rewarding effects of ethanol^87,88^. In our study, rats were treated acutely post-TBI with JZL184, and alcohol consumption testing occurred days later. Recently, our lab showed that systemic JZL184 treatment reduces the progressive-ratio breakpoint for alcohol in adult male rats subjected to a single LFP TBI^18^. Future work should test other parameters of eCB degradation enzyme inhibitors (drug, dose, repeated dosing, dosing time point, etc.) to fully characterize the effects of this drug class on alcohol-related behaviors.

We also tested rats for anxiety-like behavior 6 and 8 days after the last injury/treatment combination. It is noteworthy that most studies that report anxiolytic effects of MAGL inhibition tested animals acutely (10-120 minutes) after single^65,67,89,90^ or repeated drug treatment^69^. Here, we hypothesized that repeated systemic JZL184 treatment would rescue TBI-induced increases in anxiety-like behavior, but in fact we observed the opposite: repeated systemic treatment with JZL184 increased anxiety-like behavior in Sham animals that resembled and occluded rmTBI effects. We speculate that this may reflect a sort of “withdrawal syndrome” from repeated boosting of eCB levels via repeated MAGL inhibition, similar to what has been previously shown after prolonged pharmacological inhibition or genetic inactivation of MAGL^91,92^, but this hypothesis requires further testing. Furthermore, it is known that low doses of CB1 receptor agonists produce anxiolytic-like effects whereas high doses produce anxiogenic-like effects^67,90^. It is possible that our dosing regimen mimicked the effects of a high dose of CB1 receptor agonists, but prior work showed anxiolytic-like effects of high JZL184 doses (32mg/kg and 40mg/kg i.p) during withdrawal from methamphetamine self-administration in rats^93^. Again, it must be remembered that we administered the drug days before anxiety-like behavior testing, which differs from most prior work. Follow-up studies should measure eCB levels because chronic treatment with JZL184 (40mg/kg) increases not only levels of 2-AG but also AEA^91^, and other studies reported that simultaneous inhibition of MAGL and FAAH does not prevent stress-induced anxiety-like behavior^90^.

Based on prior literature and the quantities of alcohol consumed by animals in these experiments, it is unlikely that animals achieve physical dependence on alcohol as other models have done (e.g., vapor exposure)^94^, and that they were equally unlikely to exhibit overt withdrawal effects. That said, although we did not explicitly test it here, we cannot rule out the possibility that the forced periods without alcohol contribute to our measured outcomes.

In summary, these studies demonstrate that repeated mild TBI increases alcohol consumption in adult females and increases anxiety-like behavior in males one-week post-TBI. Sub-chronic MAGL inhibition with repeated systemic JZL184 treatments 30 min post-injury did not reduce post-TBI alcohol consumption in female rats and did not rescue post-TBI increases in anxiety-like behavior in male rats. In fact, repeated systemic JZL184 treatment mimicked and occluded TBI-induced increases in anxiety-like behavior in male rats. Future work will expand on this work by exploring other cannabinoid drugs and dosing regimens on these outcome measures after single or repeated TBIs of varying types and intensities.

## Author’s Contribution Statement

AJ-S was responsible for study design, performing experiments and collecting data, data analysis, writing/editing the manuscript. PEM was responsible for conceptualization, study design, writing/reviewing/editing the manuscript, and providing funding for the work. NWG was responsible for conceptualization, study design, writing/reviewing/editing the manuscript, and providing funding for the work.

## References

1. Dewan, MC, Rattani A, Gupta S. et al. Estimating the global incidence of traumatic brain injury. J Neurosurg. 2018; Apr 1:1-18. doi:10.3171/2017.10.JNS17352.

2. Blennow K, Brody DL, Kochanek PM, et al. Traumatic brain injuries. Nat Rev Dis Primer 2016;2:1–19. doi:10.1038/nrdp.2016.84.

3. Greco T, Ferguson L, Giza C, et al. Mechanisms underlying vulnerabilities after repeat mild traumatic brain injuries. Exp Neurol 2019;317:206–213. doi:10.1016/j.expneurol.2019.01.012.

4. Koponen S, Taiminen T, Hiekkanen H, et al. Axis I and II psychiatric disorders in patients with traumatic brain injury: a 12-month follow-up study. Brain Inj 2011;25(11):1029–34. doi:10.3109/02699052.2011.607783.

5. Vaishnavi S, Rao V, Fann JR. Neuropsychiatric Problems After Traumatic Brain Injury: Unraveling the Silent Epidemic. Psychosomatics 2009;50(3):198–205. doi:10.1176/appi.psy.50.3.198

6. Covassin T, Moran R, Elbin RJ. Sex Differences in Reported Concussion Injury Rates and Time Loss From Participation: An Update of the National Collegiate Athletic Association Injury Surveillance Program From 2004–2005 Through 2008–2009. J Athl Train 2016;51(3):189–194. doi:10.4085/1062-6050-51.3.05.

7. Lincoln AE, Caswell SV, Almquist JL, et al. Trends in concussion incidence in high school sports: a prospective 11-year study. Am J Sports Med 2011;39(5):958–963; doi: 10.1177/0363546510392326.

8. Marar M, McIlvain NM, Fields SK, et al. Epidemiology of concussions among United States high school athletes in 20 sports. Am J Sports Med 2012;40(4):747–755; doi: 10.1177/0363546511435626.

9. Colvin AC, Mullen J, Lovell MR, et al. The role of concussion history and gender in recovery from soccer-related concussion. Am J Sports Med 2009;37(9):1699–1704; doi: 10.1177/0363546509332497.

10. Wright DK, O’Brien TJ, Shultz SR, et al. Sex matters: repetitive mild traumatic brain injury in adolescent rats. Ann Clin Transl Neurol 2017;4(9):640–654; doi: 10.1002/acn3.441.

11. Alcohol Facts and Statistics | National Institute on Alcohol Abuse and Alcoholism (NIAAA). 2022. Available from: https://www.niaaa.nih.gov/publications/brochures-and-fact-sheets/alcohol-facts-and-statistics [Last accessed: 11/8/2022].

12. Savola O, Niemelä O, Hillbom M. Alcohol intake and the pattern of trauma in young adults and working aged people admitted after trauma. Alcohol Alcohol 2005;40(4):269–273; doi: 10.1093/alcalc/agh159.

13. Whelan-Goodinson R, Ponsford J, Johnston L, et al. Psychiatric disorders following traumatic brain injury: Their nature and frequency. J Head Trauma Rehabil 2009;24(5):324– 332; doi: 10.1097/HTR.0b013e3181a712aa.

14. Bombardier CH, Temkin NR, Machamer J, et al. The natural history of drinking and alcohol-related problems after traumatic brain injury. Arch Phys Med Rehabil 2003;84(2):185–91; doi: 10.1053/apmr.2003.50002.

15. Ponsford J, Whelan-Goodinson R, Bahar-Fuchs A. Alcohol and drug use following traumatic brain injury: A prospective study. Brain Inj 2007;21(13–14):1385–1392; doi: 10.1080/02699050701796960.

16. Beaulieu-Bonneau S, St-Onge F, Blackburn MC, et al. Alcohol and Drug Use before and during the First Year after Traumatic Brain Injury. J Head Trauma Rehabil 2018;33(3):E51– E60; doi: 10.1097/HTR.0000000000000341.

17. Roozenbeek B, Maas AIR, Menon DK. Changing patterns in the epidemiology of traumatic brain injury. Nat Rev Neurol 2013;9(4):231–236; doi: 10.1038/nrneurol.2013.22.

18. Fucich EA, Mayeux JP, McGinn MA, et al. A Novel Role for the Endocannabinoid System in Ameliorating Motivation for Alcohol Drinking and Negative Behavioral Affect after Traumatic Brain Injury in Rats. J Neurotrauma 2019;36(11):1847–1855; doi: 10.1089/neu.2018.5854.

19. Karelina K, Nicholson S, Weil ZM. Minocycline blocks traumatic brain injury-induced alcohol consumption and nucleus accumbens inflammation in adolescent male mice. Brain Behav Immun 2018;69:532–539; doi: 10.1016/j.bbi.2018.01.012.

20. Lowing JL, Susick LL, Caruso JP, et al. Experimental traumatic brain injury alters ethanol consumption and sensitivity. J Neurotrauma 2014;31(20):1700–10; doi: 10.1089/neu.2013.3286.

21. Mayeux JP, Teng SX, Katz PS, et al. Traumatic brain injury induces neuroinflammation and neuronal degeneration that is associated with escalated alcohol self-administration in rats. Behav Brain Res 2015;279:22–30; doi: 10.1016/j.bbr.2014.10.053.

22. Weil ZM, Karelina K, Gaier KR, et al. Juvenile Traumatic Brain Injury Increases Alcohol Consumption and Reward in Female Mice. J Neurotrauma 2016;33(9):895–903; doi: 10.1089/neu.2015.3953.

23. Lim YW, Meyer NP, Shah AS, et al. Voluntary Alcohol Intake following Blast Exposure in a Rat Model of Mild Traumatic Brain Injury. PloS One 2015;10(4):e0125130; doi: 10.1371/journal.pone.0125130.

24. Stielper ZF, Fucich EA, Middleton J, et al. Traumatic brain injury (TBI) and alcohol drinking alter basolateral amygdala (BLA) endocannabinoids in female rats. J Neurotrauma 2020; doi: 10.1089/neu.2020.7175.

25. Ceylan-Isik AF, McBride SM, Ren J. Sex Difference in Alcoholism: Who is at a Greater Risk for Development of Alcoholic Complication? Life Sci 2010;87(5–6):133–138; doi: 10.1016/j.lfs.2010.06.002.

26. Castaño-Monsalve B, Bernabeu-Guitart M, López R, et al. Alcohol and drug use disorders in patients with traumatic brain injury: neurobehavioral consequences and caregiver burden. Rev Neurol 2013;56(7):363–369.

27. Ilie G, Mann RE, Boak A, et al. Suicidality, bullying and other conduct and mental health correlates of traumatic brain injury in adolescents. PloS One 2014;9(4):e94936; doi: 10.1371/journal.pone.0094936.

28. Vaaramo K, Puljula J, Tetri S, et al. Head trauma sustained under the influence of alcohol is a predictor for future traumatic brain injury: A long-term follow-up study. Eur J Neurol 2014;21(2):293–298; doi: 10.1111/ene.12302.

29. Winqvist S, Jokelainen J, Luukinen H, et al. Adolescents’ drinking habits predict later occurrence of traumatic brain injury: 35-year follow-up of the northern Finland 1966 birth cohort. J Adolesc Health Off Publ Soc Adolesc Med 2006;39(2):275.e1–7; doi: 10.1016/j.jadohealth.2005.12.019.

30. Winqvist S, Luukinen H, Jokelainen J, et al. Recurrent traumatic brain injury is predicted by the index injury occurring under the influence of alcohol. Brain Inj 2008;22(10):780–785; doi: 10.1080/02699050802339397.

31. Xu T, Yu X, Ou S, et al. Risk factors for posttraumatic epilepsy: A systematic review and meta-analysis. Epilepsy Behav EB 2017;67:1–6; doi: 10.1016/j.yebeh.2016.10.026.

32. Hwang J, Adamson C, Butler D, et al. Enhancement of endocannabinoid signaling by fatty acid amide hydrolase inhibition: A neuroprotective therapeutic modality. Life Sci 2010;86(15–16):615–623; doi: 10.1016/j.lfs.2009.06.003.

33. Tchantchou F, Zhang Y. Selective inhibition of alpha/beta-hydrolase domain 6 attenuates neurodegeneration, alleviates blood brain barrier breakdown, and improves functional recovery in a mouse model of traumatic brain injury. J Neurotrauma 2013;30(7):565–579; doi: 10.1089/neu.2012.2647.

34. Blankman JL, Simon GM, Cravatt BF. A comprehensive profile of brain enzymes that hydrolyze the endocannabinoid 2-arachidonoylglycerol. Chem Biol 2007;14(12):1347–1356; doi: 10.1016/j.chembiol.2007.11.006.

35. Mayeux J, Katz P, Edwards S, et al. Inhibition of Endocannabinoid Degradation Improves Outcomes from Mild Traumatic Brain Injury: A Mechanistic Role for Synaptic Hyperexcitability. J Neurotrauma 2017;34(2):436–443; doi: 10.1089/neu.2016.4452.

36. Zhang J, Teng Z, Song Y, et al. Inhibition of monoacylglycerol lipase prevents chronic traumatic encephalopathy-like neuropathology in a mouse model of repetitive mild closed head injury. J Cereb Blood Flow Metab Off J Int Soc Cereb Blood Flow Metab 2015;35(3):443–453; doi: 10.1038/jcbfm.2014.216.

37. Katz PS, Sulzer JK, Impastato RA, et al. Endocannabinoid degradation inhibition improves neurobehavioral function, blood-brain barrier integrity, and neuroinflammation following mild traumatic brain injury. J Neurotrauma 2015;32(5):297–306; doi: 10.1089/neu.2014.3508.

38. Hawley LA, Ketchum JM, Morey C, et al. Cannabis Use in Individuals With Spinal Cord Injury or Moderate to Severe Traumatic Brain Injury in Colorado. Arch Phys Med Rehabil 2018;99(8):1584–1590; doi: 10.1016/j.apmr.2018.02.003.

39. Bodnar RJ. Endogenous Opiates and Behavior: 2014. Elsevier Inc.; 2016.; doi: 10.1016/j.peptides.2015.10.009.

40. Long JZ, Li W, Booker L, et al. Selective blockade of 2-arachidonoylglycerol hydrolysis produces cannabinoid behavioral effects. Nat Chem Biol 2009;5(1):37–44; doi: 10.1038/nchembio.129.

41. Teng SX, Molina PE. Acute alcohol intoxication prolongs neuroinflammation without exacerbating neurobehavioral dysfunction following mild traumatic brain injury. J Neurotrauma 2014;31(4):378–386; doi: 10.1089/neu.2013.3093.

42. Fujita M, Wei EP, Povlishock JT. Intensity- and interval-specific repetitive traumatic brain injury can evoke both axonal and microvascular damage. J Neurotrauma 2012;29(12):2172–2180; doi: 10.1089/neu.2012.2357.

43. Huang L, Coats JS, Mohd-Yusof A, et al. Tissue vulnerability is increased following repetitive mild traumatic brain injury in the rat. Brain Res 2013;1499:109–120; doi: 10.1016/j.brainres.2012.12.038.

44. Laurer HL, Bareyre FM, Lee VM, et al. Mild head injury increasing the brain’s vulnerability to a second concussive impact. J Neurosurg 2001;95(5):859–870; doi: 10.3171/jns.2001.95.5.0859.

45. Longhi L, Saatman KE, Fujimoto S, et al. Temporal window of vulnerability to repetitive experimental concussive brain injury. Neurosurgery 2005;56(2):364–374; discussion 364-374; doi: 10.1227/01.neu.0000149008.73513.44.

46. Mouzon B, Chaytow H, Crynen G, et al. Repetitive mild traumatic brain injury in a mouse model produces learning and memory deficits accompanied by histological changes. J Neurotrauma 2012;29(18):2761–2773; doi: 10.1089/neu.2012.2498.

47. Mouzon BC, Bachmeier C, Ferro A, et al. Chronic neuropathological and neurobehavioral changes in a repetitive mild traumatic brain injury model. Ann Neurol 2014;75(2):241–254; doi: 10.1002/ana.24064.

48. Prins ML, Alexander D, Giza CC, et al. Repeated mild traumatic brain injury: mechanisms of cerebral vulnerability. J Neurotrauma 2013;30(1):30–38; doi: 10.1089/neu.2012.2399.

49. Management of Concussion/mTBI Working Group. VA/DoD Clinical Practice Guideline for Management of Concussion/Mild Traumatic Brain Injury. J Rehabil Res Dev 2009;46(6):CP1-68.

50. Dewitt DS, Perez-Polo R, Hulsebosch CE, et al. Challenges in the development of rodent models of mild traumatic brain injury. J Neurotrauma 2013;30(9):688–701; doi: 10.1089/neu.2012.2349.

51. Cernak I. Animal models of head trauma. NeuroRx 2005;2(3):410–422; doi: 10.1602/neurorx.2.3.410.

52. Gronwall D, Wrightson P. Cumulative effect of concussion. Lancet Lond Engl 1975;2(7943):995–997; doi: 10.1016/s0140-6736(75)90288-3.

53. Rabadi MH, Jordan BD. The cumulative effect of repetitive concussion in sports. Clin J Sport Med Off J Can Acad Sport Med 2001;11(3):194–198; doi: 10.1097/00042752-200107000-00011.

54. Holgate JY, Shariff M, Mu EWH, et al. A Rat Drinking in the Dark Model for Studying Ethanol and Sucrose Consumption. Front Behav Neurosci 2017;11:29; doi: 10.3389/fnbeh.2017.00029.

55. Simms JA, Steensland P, Medina B, et al. Intermittent access to 20% ethanol induces high ethanol consumption in Long-Evans and Wistar rats. Alcohol Clin Exp Res 2008;32(10):1816–1823; doi: 10.1111/j.1530-0277.2008.00753.x.

56. Schindler AG, Baskin B, Juarez B, et al. Repetitive blast mild traumatic brain injury increases ethanol sensitivity in male mice and risky drinking behavior in male combat veterans. Alcohol Clin Exp Res 2021;45(5):1051–1064; doi: 10.1111/acer.14605.

57. Hoffman J, Yu J, Kirstein C, et al. Combined Effects of Repetitive Mild Traumatic Brain Injury and Alcohol Drinking on the Neuroinflammatory Cytokine Response and Cognitive Behavioral Outcomes. Brain Sci 2020;10(11):E876; doi: 10.3390/brainsci10110876.

58. Gupte R, Brooks W, Vukas R, et al. Sex Differences in Traumatic Brain Injury: What We Know and What We Should Know. J Neurotrauma 2019;36(22):3063–3091; doi: 10.1089/neu.2018.6171.

59. Späni CB, Braun DJ, Van Eldik LJ. Sex-related responses after traumatic brain injury: considerations for preclinical modeling. Front Neuroendocrinol 2018;50:52–66; doi: 10.1016/j.yfrne.2018.03.006.

60. Winston CN, Noël A, Neustadtl A, et al. Dendritic Spine Loss and Chronic White Matter Inflammation in a Mouse Model of Highly Repetitive Head Trauma. Am J Pathol 2016;186(3):552–567; doi: 10.1016/j.ajpath.2015.11.006.

61. Petraglia AL, Plog BA, Dayawansa S, et al. The spectrum of neurobehavioral sequelae after repetitive mild traumatic brain injury: a novel mouse model of chronic traumatic encephalopathy. J Neurotrauma 2014;31(13):1211–1224; doi: 10.1089/neu.2013.3255.

62. Christensen J, Eyolfson E, Salberg S, et al. When Two Wrongs Make a Right: The Effect of Acute and Chronic Binge Drinking on Traumatic Brain Injury Outcomes in Young Adult Female Rats. J Neurotrauma 2020;37(2):273–285; doi: 10.1089/neu.2019.6656.

63. Chandrasekar A, Aksan B, Heuvel FO, et al. Neuroprotective effect of acute ethanol intoxication in TBI is associated to the hierarchical modulation of early transcriptional responses. Exp Neurol 2018;302:34–45; doi: 10.1016/j.expneurol.2017.12.017.

64. Brown KJ, Grunberg NE. Effects of housing on male and female rats: crowding stresses male but calm females. Physiol Behav 1995;58(6):1085–1089; doi: 10.1016/0031-9384(95)02043-8.

65. Busquets-Garcia A, Puighermanal E, Pastor A, et al. Differential role of anandamide and 2-arachidonoylglycerol in memory and anxiety-like responses. Biol Psychiatry 2011;70(5):479–486; doi: 10.1016/j.biopsych.2011.04.022.

66. Kathuria S, Gaetani S, Fegley D, et al. Modulation of anxiety through blockade of anandamide hydrolysis. Nat Med 2003;9(1):76–81; doi: 10.1038/nm803.

67. Patel S, Hillard CJ. Pharmacological Evaluation of Cannabinoid Receptor Ligands in a Mouse Model of Anxiety: Further Evidence for an Anxiolytic Role for Endogenous Cannabinoid Signaling. J Pharmacol Exp Ther 2006;318(1):304–311; doi: 10.1124/jpet.106.101287.

68. Patel S, Hill MN, Cheer JF, et al. The endocannabinoid system as a target for novel anxiolytic drugs. Neurosci Biobehav Rev 2017;76(Pt A):56–66; doi: 10.1016/j.neubiorev.2016.12.033.

69. Sciolino NR, Zhou W, Hohmann AG. Enhancement of endocannabinoid signaling with JZL184, an inhibitor of the 2-arachidonoylglycerol hydrolyzing enzyme monoacylglycerol lipase, produces anxiolytic effects under conditions of high environmental aversiveness in rats. Pharmacol Res 2011;64(3):226–234; doi: 10.1016/j.phrs.2011.04.010.

70. Haller J, Bakos N, Szirmay M, et al. The effects of genetic and pharmacological blockade of the CB1 cannabinoid receptor on anxiety. Eur J Neurosci 2002;16(7):1395–1398; doi: 10.1046/j.1460-9568.2002.02192.x.

71. Haller J, Varga B, Ledent C, et al. CB1 cannabinoid receptors mediate anxiolytic effects: convergent genetic and pharmacological evidence with CB1-specific agents. Behav Pharmacol 2004;15(4):299–304; doi: 10.1097/01.fbp.0000135704.56422.40.

72. Martin M, Ledent C, Parmentier M, et al. Involvement of CB1 cannabinoid receptors in emotional behaviour. Psychopharmacology (Berl) 2002;159(4):379–387; doi: 10.1007/s00213-001-0946-5.

73. Colombo G, Serra S, Brunetti G, et al. Stimulation of voluntary ethanol intake by cannabinoid receptor agonists in ethanol-preferring sP rats. Psychopharmacology (Berl) 2002;159(2):181–187; doi: 10.1007/s002130100887.

74. Gallate JE, Saharov T, Mallet PE, et al. Increased motivation for beer in rats following administration of a cannabinoid CB1 receptor agonist. Eur J Pharmacol 1999;370(3):233– 240; doi: 10.1016/s0014-2999(99)00170-3.

75. Getachew B, Hauser SR, Dhaher R, et al. CB1 receptors regulate alcohol-seeking behavior and alcohol self-administration of alcohol-preferring (P) rats. Pharmacol Biochem Behav 2011;97(4):669–675; doi: 10.1016/j.pbb.2010.11.006.

76. Alén F, Santos A, Moreno-Sanz G, et al. Cannabinoid-induced increase in relapse-like drinking is prevented by the blockade of the glycine-binding site of N-methyl-D-aspartate receptors. Neuroscience 2009;158(2):465–473; doi: 10.1016/j.neuroscience.2008.10.002.

77. Economidou D, Mattioli L, Cifani C, et al. Effect of the cannabinoid CB1 receptor antagonist SR-141716A on ethanol self-administration and ethanol-seeking behaviour in rats. Psychopharmacology (Berl) 2006;183(4):394–403; doi: 10.1007/s00213-005-0199-9.

78. Femenía T, García-Gutiérrez MS, Manzanares J. CB1 receptor blockade decreases ethanol intake and associated neurochemical changes in fawn-hooded rats. Alcohol Clin Exp Res 2010;34(1):131–141; doi: 10.1111/j.1530-0277.2009.01074.x.

79. Freedland CS, Sharpe AL, Samson HH, et al. Effects of SR141716A on ethanol and sucrose self-administration. Alcohol Clin Exp Res 2001;25(2):277–282.

80. Arnone M, Maruani J, Chaperon F, et al. Selective inhibition of sucrose and ethanol intake by SR 141716, an antagonist of central cannabinoid (CB1) receptors. Psychopharmacology (Berl) 1997;132(1):104–106; doi: 10.1007/s002130050326.

81. Cippitelli A, Bilbao A, Hansson AC, et al. Cannabinoid CB1 receptor antagonism reduces conditioned reinstatement of ethanol-seeking behavior in rats. Eur J Neurosci 2005;21(8):2243–2251; doi: 10.1111/j.1460-9568.2005.04056.x.

82. Colombo G, Vacca G, Serra S, et al. Suppressing effect of the cannabinoid CB1 receptor antagonist, SR 141716, on alcohol’s motivational properties in alcohol-preferring rats. Eur J Pharmacol 2004;498(1–3):119–123; doi: 10.1016/j.ejphar.2004.07.069.

83. Gessa GL, Serra S, Vacca G, et al. Suppressing effect of the cannabinoid CB1 receptor antagonist, SR147778, on alcohol intake and motivational properties of alcohol in alcohol-preferring sP rats. Alcohol Alcohol 2005;40(1):46–53; doi: 10.1093/alcalc/agh114.

84. Lallemand F, Soubrié PH, De Witte PH. Effects of CB1 cannabinoid receptor blockade on ethanol preference after chronic ethanol administration. Alcohol Clin Exp Res 2001;25(9):1317–1323.

85. Rodríguez de Fonseca F, Roberts AJ, Bilbao A, et al. Cannabinoid receptor antagonist SR141716A decreases operant ethanol self administration in rats exposed to ethanol-vapor chambers. Zhongguo Yao Li Xue Bao 1999;20(12):1109–1114.

86. Rácz I, Markert A, Mauer D, et al. Long-term ethanol effects on acute stress responses: modulation by dynorphin. Addict Biol 2013;18(4):678–688; doi: 10.1111/j.1369-1600.2012.00494.x.

87. Houchi H, Babovic D, Pierrefiche O, et al. CB1 receptor knockout mice display reduced ethanol-induced conditioned place preference and increased striatal dopamine D2 receptors. Neuropsychopharmacol Off Publ Am Coll Neuropsychopharmacol 2005;30(2):339–349; doi: 10.1038/sj.npp.1300568.

88. Naassila M, Pierrefiche O, Ledent C, et al. Decreased alcohol self-administration and increased alcohol sensitivity and withdrawal in CB1 receptor knockout mice. Neuropharmacology 2004;46(2):243–253; doi: 10.1016/j.neuropharm.2003.09.002.

89. Almeida-Santos AF, Gobira PH, Rosa LC, et al. Modulation of anxiety-like behavior by the endocannabinoid 2-arachidonoylglycerol (2-AG) in the dorsolateral periaqueductal gray. Behav Brain Res 2013;252:10–17; doi: 10.1016/j.bbr.2013.05.027.

90. Bedse G, Bluett RJ, Patrick TA, et al. Therapeutic endocannabinoid augmentation for mood and anxiety disorders: comparative profiling of FAAH, MAGL and dual inhibitors. Transl Psychiatry 2018;8(1):92; doi: 10.1038/s41398-018-0141-7.

91. Schlosburg JE, Blankman JL, Long JZ, et al. Chronic monoacylglycerol lipase blockade causes functional antagonism of the endocannabinoid system. Nat Neurosci 2010;13(9):1113–1119; doi: 10.1038/nn.2616.

92. Schlosburg JE, Kinsey SG, Ignatowska-Jankowska B, et al. Prolonged monoacylglycerol lipase blockade causes equivalent cannabinoid receptor type 1 receptor-mediated adaptations in fatty acid amide hydrolase wild-type and knockout mice. J Pharmacol Exp Ther 2014;350(2):196–204; doi: 10.1124/jpet.114.212753.

93. Nawata Y, Yamaguchi T, Fukumori R, et al. Inhibition of Monoacylglycerol Lipase Reduces the Reinstatement of Methamphetamine-Seeking and Anxiety-Like Behaviors in Methamphetamine Self-Administered Rats. Int J Neuropsychopharmacol 2019;22(2):165– 172; doi: 10.1093/ijnp/pyy086.

94. Avegno EM, Gilpin NW. Inducing Alcohol Dependence in Rats Using Chronic Intermittent Exposure to Alcohol Vapor. Bio-Protoc 2019;9(9):e3222; doi: 10.21769/BioProtoc.3222.

